# Immunomodulatory activity of *Ganoderma lucidum* immunomodulatory protein *via* PI3K/Akt and MAPK signaling pathways in macrophage RAW264.7 cells

**DOI:** 10.1101/499871

**Authors:** Qi-Zhang Li, Yu-Zhou Chang, Liu-Ding-Ji Li, Xiao-Yu Du, Xiao-Hui Bai, Zhu-Mei He, Lei Chen, Xuan-Wei Zhou

## Abstract

*Ganoderma lucidum*, a traditional edible and medicinal fungus, holds an important status in health care systems in China and other Asian countries. Fungal immunomodulatory protein (FIP), one of the active ingredients isolated from *G. lucidum*, is a class of naturally occurring proteins and possesses potential biological functions. This study was conducted to explore the molecular mechanism of its immunomodulatory potency in immune responses of macrophages. *In vitro* assays of biological activity indicated that rFIP-glu significantly activated macrophage RAW264.7 cells, and possessed the ability of pro-and anti-inflammation the cells. RNA sequencing analysis showed that macrophage activation involved Toll-like receptors and mitogen-activated protein kinases pathways. Furthermore, qRT-PCR indicated that phosphoinositide 3 kinase inhibitor LY294002 blocked the mRNA levels of MCP-1, MEK1/2 inhibitor U0126 reduced the mRNA levels of TNF-α and MCP-1, and JNK inhibitor SP600125 prevented the up-regulation of iNOS mRNA in the rFIP-glu-induced cells. FIP-glu mediated these inflammatory effects not through a general pathway, instead through a different pathway for different inflammatory mediator. These data indicate the possibility that rFIP-glu has an important immune-regulation function and thus has potential therapeutic uses.

---

*Ganoderma lucidum*, a traditional Chinese medicinal mushroom, is a species in the genus of *Ganoderma* with numerous pharmacological effects such as improving immune function, antitumor, antioxidant, reducing cardiovascular and cerebrovascular diseases and heart diseases caused by body oxidation (1). Among more than 400 different bioactive compounds isolated from *G. lucidum*, fungal immunomodulatory protein (FIP) is an important bioactive component with immune regulating activity and is one of the most promising active ingredients developed by modern biotechnologies (2). FIP is a small protein with similar structure and immune-regulatory activity to phytohemagglutinin and immunoglobulins. Since the first FIP (designated as Lingzhi-8 or LZ-8) was isolated from *G. lucidum* mycelia, dozens of FIPs have been isolated and identified from different fungous species in recent years (3–6). FIPs have immunomodulatory functions and play an important role in anti-tumor, anti-allergy, anti-transplant rejection, *etc*., which implies a promising application for medicinal use (7). For example, FIPs suppress tumors by inhibition of telomerase activity *via* decrease of *hTERT* promoter activity and translocation of its protein (8–10). FIP-gmi induce apoptosis *via* β-catenin inhibition in lung cancer cells (11). FIP-fve has anti-inflammatory effects on OVA-induced airway inflammation and reduces airway remodeling by suppressing IL-17 (12). In spite of a few studies about anti-tumor (13, 14) and immunomodulatory effects (15–17), the mechanism of these activities remain unclear. The relationship between the activation of these proteins, downstream cytokine expression and physiological function represents an active line of investigation.

Macrophages, belonging to a group of mononuclear phagocytes, play vital roles in processes of the immune response and are strategically positioned throughout the body tissues (18). They possess functions of phagocytosis, antigen presentation and production of cytokines, thereby initiating immune response (19). Following activation, macrophages can release a wide array of pro-or anti-inflammatory cytokines, which further activate fellow immune cells (20). Depending on these signals, macrophages have been typed classically activated (pro-inflammation, M1) and alternatively activated (anti-inflammation, M2) (21). Classically activated macrophages are elicited in response to pro-inflammatory cytokines and pathogen-associated molecular patterns (PAMPs), such as Interferon-γ (IFN-γ) and lipopolysaccharide (LPS) to promote pathogen killing and chronic inflammation. (22, 23). These macrophages produce cytotoxic and inflammatory molecules nitric oxide (NO) and reactive oxygen species (ROS), pro-inflammatory cytokines tumor necrosis factor (TNF-α), interleukin (IL)-1β and IL-6, chemokine monocyte chemoattractant protein-1 (MCP-1, or C-C motif ligand 2 (CCL-2)), *etc*. (24). However, excessive inflammatory mediators can be implicated in a number of chronic diseases, such as arthritis, colitis and asthma (25). M2 macrophages, which are typed in response to anti-inflammatory cytokines, parasitic infections and damage-associated molecular patterns (DAMPs), such as IL-4 and IL-13, play an important role in inhibition of chronic and acute inflammatory response and in tissue repair (23). M2 macrophages are able to secrete high amounts of anti-inflammatory cytokines, such as IL-10 and TGF-β (26).

Macrophages can be activated to an inflammation-promoting phenotype through members of the Toll-like receptor (TLR) family such as TLR4 (20, 27, 28). Activated TLRs induce activation of specific intracellular pathways including phosphoinositide 3 kinases (PI3K/Akt), mitogen activated protein kinases (MAPKs) and nuclear factor kappa B (NF-κB) (21, 29, 30). PI3K/Akt signaling pathway participates in macrophage polarization (31–34), while MAPKs, including extracellular signal-related kinase (ERK)-1/2, p38 and c-Jun NH2-terminal kinase (JNK), and NF-κB are classic inflammation related signals and induce the expression of pro-inflammatory mediators (35–38).

Here, we report that rFIP-glu produced in *Pichia pastoris* has the ability to induce macrophage activation and produce pro-and anti-inflammatory mediators, which may be through PI3K and MAPK pathways.

## RESULTS

### Production of rFIP-glu in *Pichia pastoris*

An expression vector pPIC9K was used for achieving rFIP-glu. To facilitate purification, a His-tag was added at the C-terminal of rFIP-glu (Fig. 1A). Following confirmation by sequencing and linearization by *Sac I*, a recombination plasmid containing nucleotide sequences of FIP-glu and His-tag was transformed into *Pichia pastoris* GS115 cells. The transformants successfully secreted recombination proteins into the media compared with the negative control (GS115 transformed with or without pPIC9K plasmid incubated with or without MeOH) after induced by MeOH for 72 h (Fig. 1B). Western blot analysis further confirmed that the secreted protein was rFIP-glu using anti-6×His tag (Fig. 1C) and anti-rFIP-glu (Fig. 1D).

**FIG 1.**
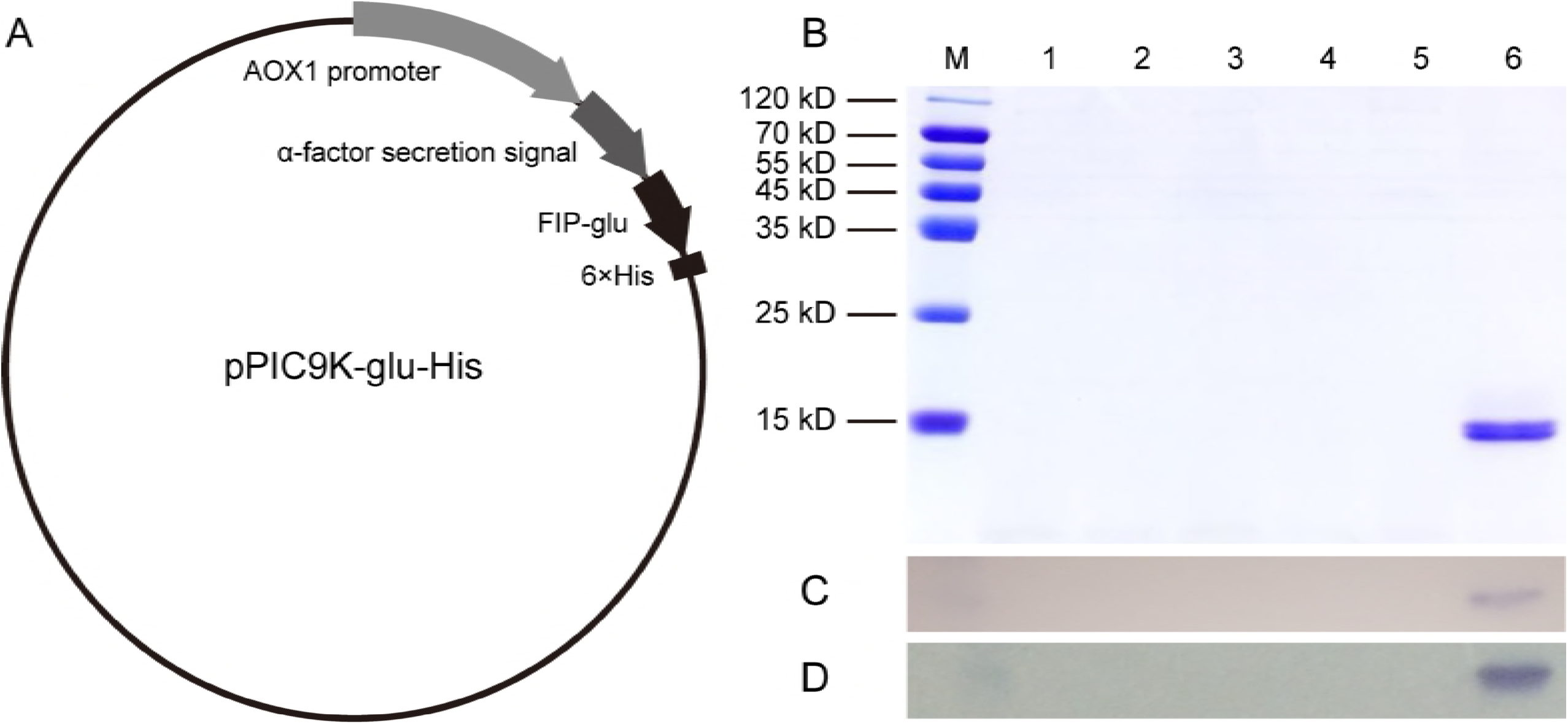
Production of rFIP-glu in *P. pastoris* GS115 cells. A, schematic map of the recombinant expression vector pPIC9K-glu-His. B, rFIP-glu detected by SDS-PAGE. C, Western blot analysis of rFIP-glu using anti-6×His tag. D, Western blot analysis of rFIP-glu using anti-rFIP-glu. Lane M: protein molecular mass marker. Lane 1, *P. pastoris* GS115 cells without being induced by MeOH. Lane 2, *P. pastoris* GS115 cells induced by MeOH. Lane 3, *P. pastoris* GS115 cells containing expression vector pPIC9K without being induced by MeOH. Lane 4, *P. pastoris* GS115 cells containing expression vector pPIC9K induced by MeOH. Lane 5, *P. pastoris* GS115 cells containing expression vector pPIC9K-glu-His without being induced by MeOH. Lane 6, *P. pastoris* GS115 cells containing expression vector pPIC9K-glu-His induced by MeOH.

### Toxicity of rFIP-glu against RAW264.7 cells

To investigate the activation effect of rFIP-glu on the RAW264.7 cells, firstly we determined whether this recombinant fungal protein possessed toxicity and measured its noncytotoxic range. As shown in Fig. 2, the proliferation of RAW265.7 cells treated with 1 and 2 μg/mL of rFIP-glu significantly increased (*p* ≤ 0.0001), relative to the control group. A culture of RAW264.7 cells incubated with 4 μg/mL of rFIP-glu resulted in no effect on cell viability and more than 90% of cells with 8 μg/mL were viable (*p* ≤ 0.01). The viability of RAW264.7 cells with more than 8 μg/mL showed a rapid decease (*p* ≤ 0.0001). Based on these results, subsequent assays were performed at 4 μg/mL or no more than 8 μg/mL. Additionally, rFIP-glu obviously influenced the morphology of RAW264.7 macrophages with or without stimulation of LPS (Fig. S1).

**FIG 2.**
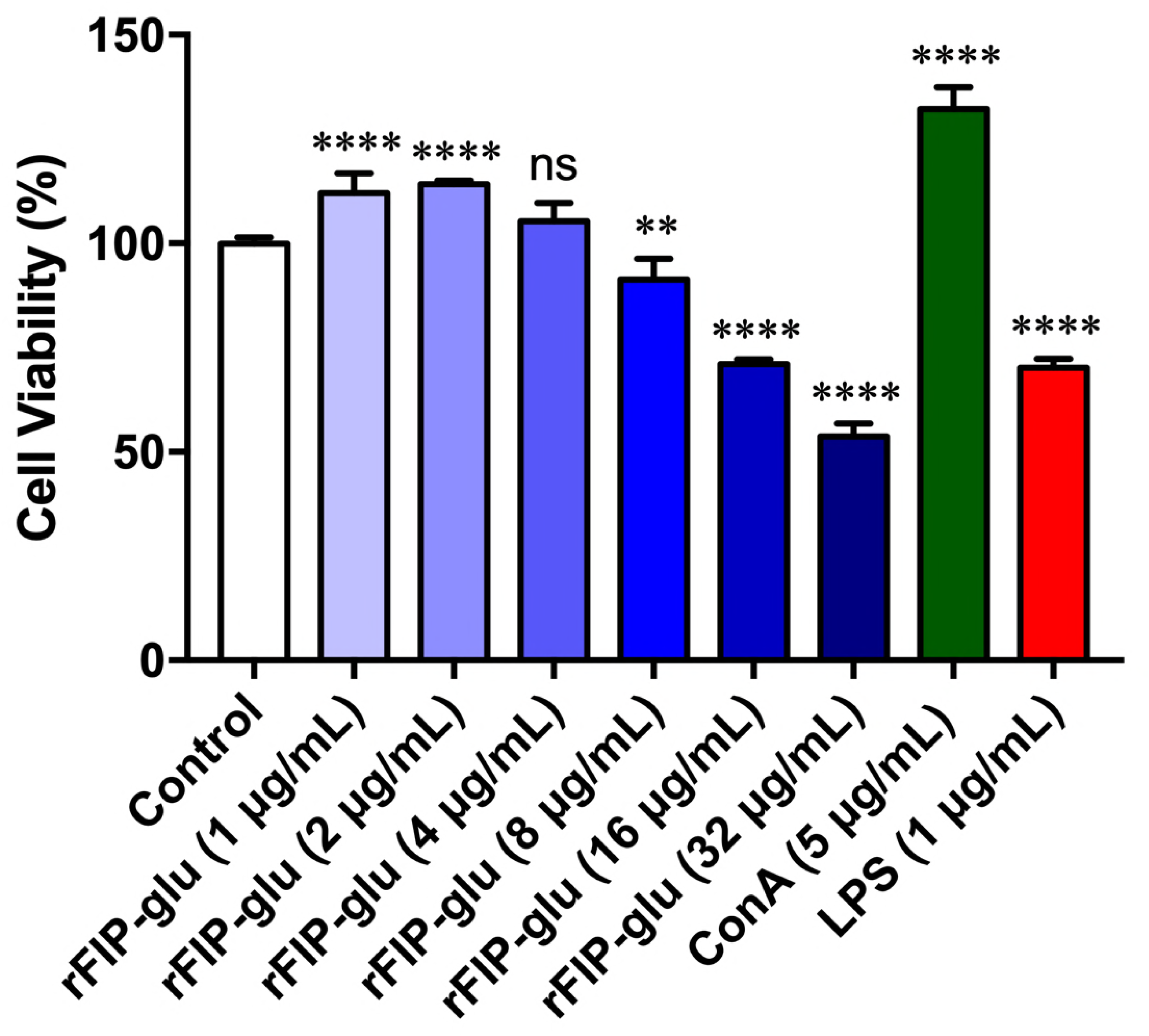
Effect of rFIP-glu on the viability of RAW264.7 cells. RAW264.7 cells were treated with different concentration of rFIP-glu (1, 2, 4, 8, 16 and 32 μg/mL) or PBS (as control), Concanavalin A (ConA) and lipopolysaccharide (LPS) for 24 h. The cells were measured by methylene blue uptake assay. Data were expressed as means ± SD (n = 5). ns, *p* > 0.05; **, *p* ≤ 0.01; ****, *p* ≤ 0.0001 versus control group.

### rFIP-glu improves phagocytosis of RAW264.7 cells

Next, phagocytic activities of RAW264.7 cells treated with rFIP-glu were examined by neutral red uptake assay. As shown in Fig. 3, when the cells were treated with noncytotoxic concentration of rFIP-glu ranged from 1 to 8 μg/mL, the phagocytosis increased and then decreased. The phagocytosis of macrophage RAW264.7 cells were significantly improved (*p* ≤ 0.05) at 2 μg/mL of rFIP-glu, but suppressed (*p* > 0.05) at 8 μg/mL. These indicate that rFIP-glu has the ability to enhance phagocytic activity of RAW 264.7 cells when the concentration is no higher than 8 μg/mL.

**FIG 3.**
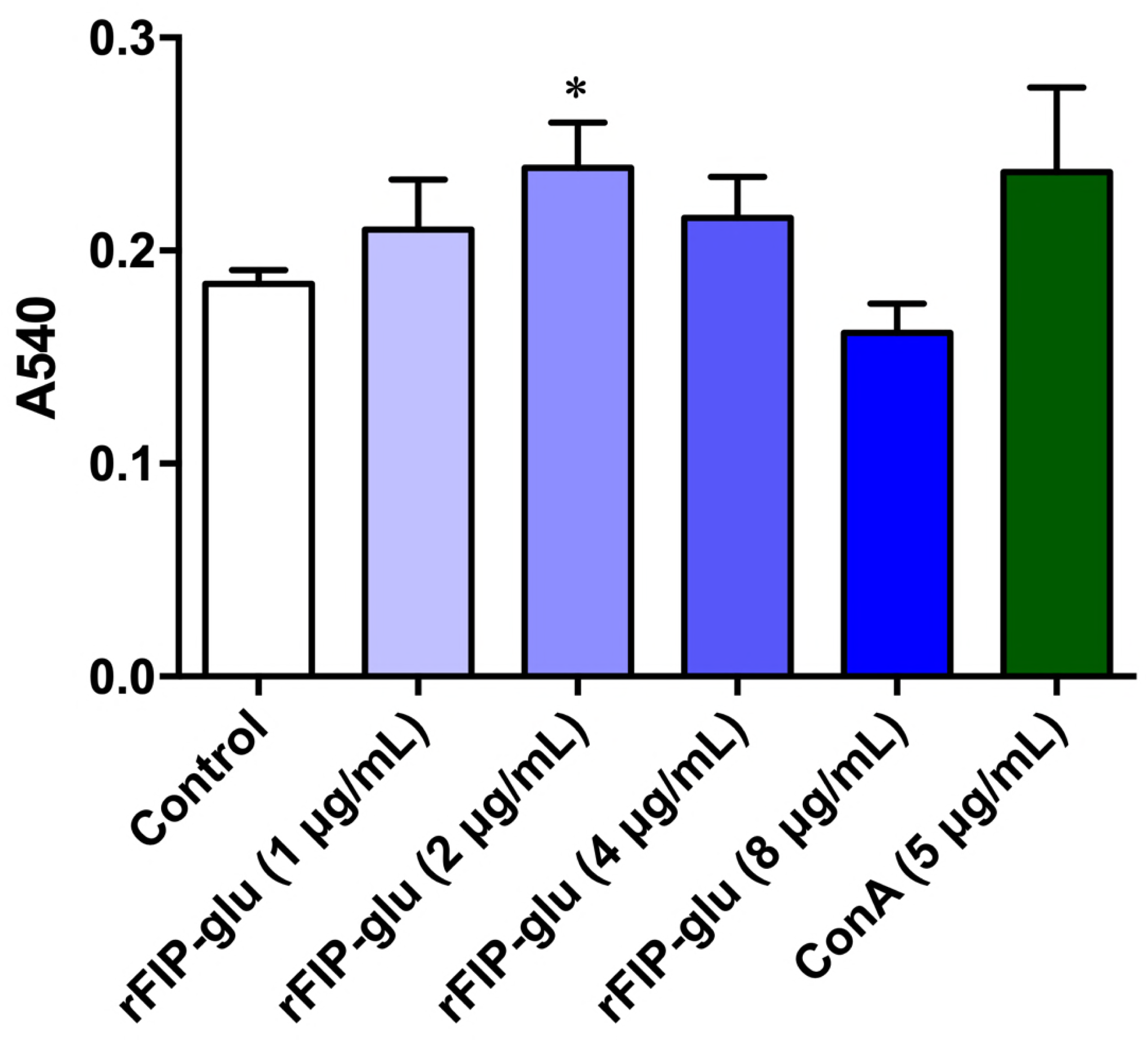
Effects of rFIP-glu on RAW264.7 cells phagocytosis. RAW264.7 cells were treated with different concentration of rFIP-glu (1, 2, 4 and 8 μg/mL). Control cells were treated with PBS only and ConA cells as positive control were treated with 5 μg/mL. The effects were assessed by neutral red uptake assay. Data were expressed as means ± SD (n = 3). *, *p* ≤ 0.05 versus control group.

### rFIP-glu regulates pro-and anti-inflammatory genes at transcriptional level in RAW264.7 cells

To further evaluate the immunostimulatory effects of rFIP-glu, we investigated whether rFIP-glu had the ability to induce mRNA levels of relevant genes contributing to the function of macrophages. These genes were determined by qRT-PCR after cells were treated with rFIP-glu (1, 2, 4 and 8 μg/mL) for 6 h (Fig. 4). Compared to control, the rFIP-glu-treated group showed a robust increase in the mRNA level of TNF-α (Fig. 4A; *p* ≤ 0.01 at 1 μg/mL; *p* ≤ 0.001 at 2, 4 and 8 μg/mL). Similarly, rFIP-glu also significantly promoted the mRNA expression of Arginase II in a concentration-dependent manner (Fig. 4B; *p* ≤ 0.001 at 1 μg/mL; *p* ≤ 0.0001 at 2, 4 and 8 μg/mL). The production of NO was measured firstly, but none was detected at 1 to 8 μg/mL of rFIP-glu (Data not shown). Alternatively, we investigated the mRNA expression of iNOS and found that it also dramatically increased in a concentration-dependent manner (Fig. 4C; *p* ≤ 0.0001 at 4 and 8 μg/mL). The mRNA level of MCP-1 (CCL-2) was also increased and peaked at 2 μg/mL (*p* ≤ 0.0001), and then decreased to no change at 8 μg/mL compared with control (Fig. 4D). rFIP-glu treatment concentration-dependently inhibited the mRNA expression levels of IL-10 (Fig. 4E; *p* ≤ 0.0001) in RAW264.7 cells. These results exhibit that rFIP-glu stimulates the immune responses by inducing pro-inflammatory mediators. In parallel, rFIP-glu inhibited the mRNA expression level of CXCL-10 (Fig. 4F; *p* ≤ 0.01 at 1 μg/mL; *p* ≤ 0.0001 at 2, 4 and 8 μg/mL). This result exhibits that rFIP-glu induces anti-inflammatory phenotype of macrophages. Additionally, there was little or no effect on IL-6 at transcriptional level (Fig. 4G) and the mRNA expression of IL-1β was not detected (Data not shown). Taken together, rFIP-glu re-polarizes macrophages by regulating of pro-and anti-inflammatory genes expression at transcriptional level in RAW264.7 cells.

**FIG 4.**
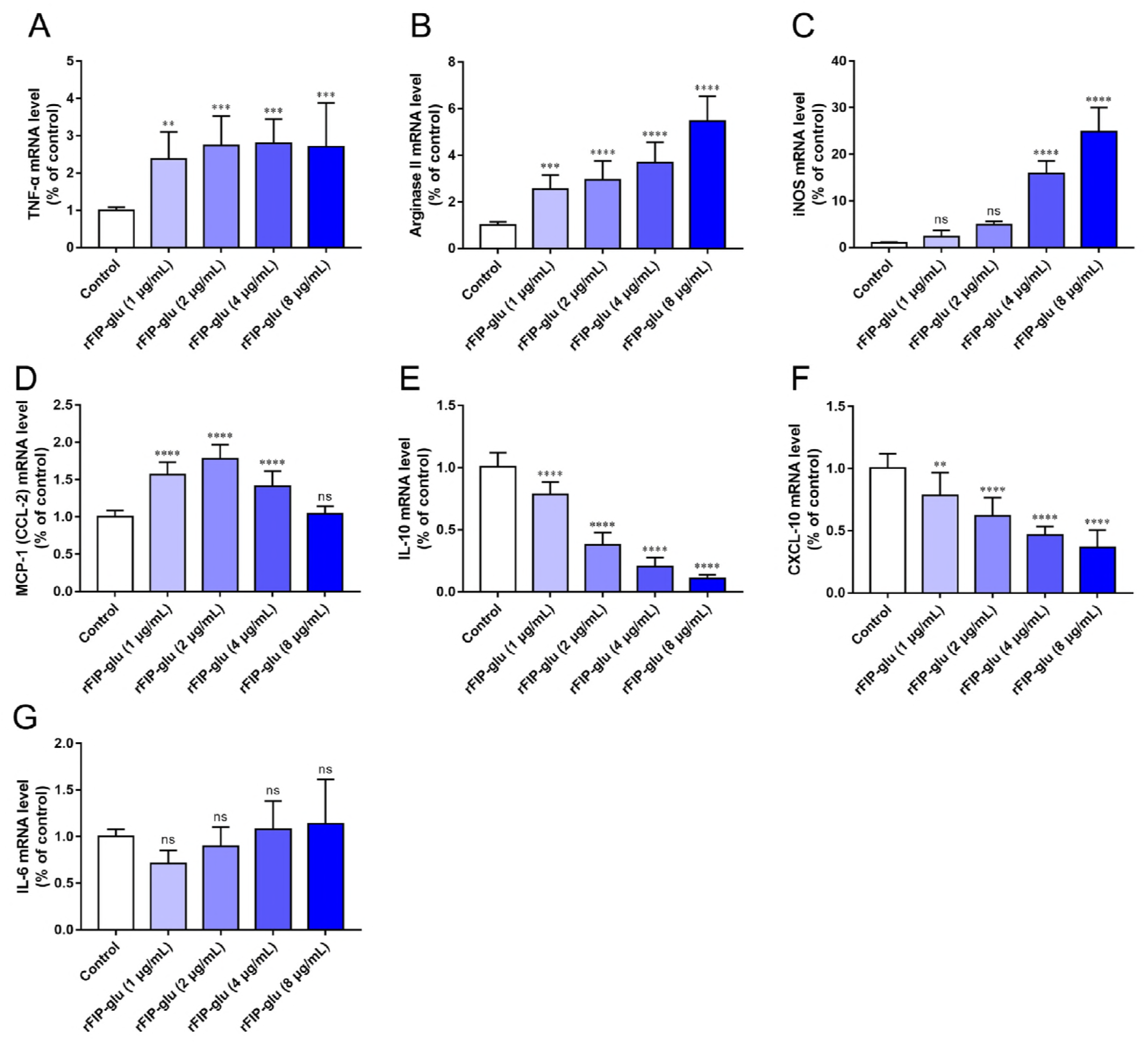
Effects of rFIP-glu on RAW264.7 cells. The cells were treated with different concentration of rFIP-glu (1, 2, 4 and 8 μg/mL) for 6 h. The mRNA expression of TNF-α (A), Arginase II (B), iNOS (C), MCP-1 (CCL-2) (D), IL-10 (E), CXCL-10 (F) and IL-6 (G) was measured by qRT-PCR. Data were expressed as means ± SD (n = 3). ns, *p* > 0.05; **, *p* ≤ 0.01; ***, *p* ≤ 0.001; ****, *p* ≤ 0.0001 versus control group.

### rFIP-glu regulates LPS-induced pro-and anti-inflammatory mediators at transcriptional level in RAW264.7 cells

In order to further determine that rFIP-glu induces or suppresses mRNA expression levels of pro-and anti-inflammatory genes, RAW264.7 cells were stimulated with 1 μg/mL LPS in the presence or absence of increasing concentration of rFIP-glu for 6 h, and then qRT-PCR analysis was performed. Compared with the normal control group, LPS treatment (1 μg/mL) significantly increased the mRNA expression of IL-1β, IL-6, IL-10, TNF-α, MCP-1 (CCL-2), CXCL-10 and Arginase II and production of NO (Fig. 5). rFIP-glu treatment (1, 2, 4 and 8 μg/mL) significantly promoted the mRNA levels of TNF-α (Fig. 5A; *p* ≤ 0.0001), MCP-1 (CCL-2) (Fig. 5B; *p* ≤ 0.0001) and Arginase II (Fig. 5C; *p* ≤ 0.0001) and concentration-dependently inhibited IL-10 expression at transcriptional level (Fig. 5D; *p* ≤ 0.001 at 2 μg/mL; *p* ≤ 0.0001 at 4 and 8 μg/mL) in LPS-stimulated RAW 264.7 cells. These results suggest that rFIP-glu promotes inflammation in LPS-induced RAW264.7. On the other hand, rFIP-glu could suppressed LPS-induced mRNA expression levels of IL-1β (Fig. 5E; *p* ≤ 0.01 at 2 μg/mL; *p* ≤ 0.0001 at 4 and 8 μg/mL), IL-6 (Fig. 5F; *p* ≤ 0.01 at 4 μg/mL; *p* ≤ 0.0001 at 8 μg/mL) and CXCL-10 (Fig. 5G; *p* ≤ 0.01 at 4 μg/mL; *p* ≤ 0.0001 at 8 μg/mL) and LPS-induced production of NO (Fig. 5H; *p* ≤ 0.01 at 2 μg/mL; *p* ≤ 0.0001 at 4 and 8 μg/mL) in a concentration-dependent manner. The results indicate that rFIP-glu suppresses the LPS-induced expression of these inflammatory mediators at the transcriptional level. Thus, rFIP-glu regulates pro-and anti-inflammatory genes expression at transcriptional level in LPS-stimulated RAW264.7 cells.

**FIG 5.**
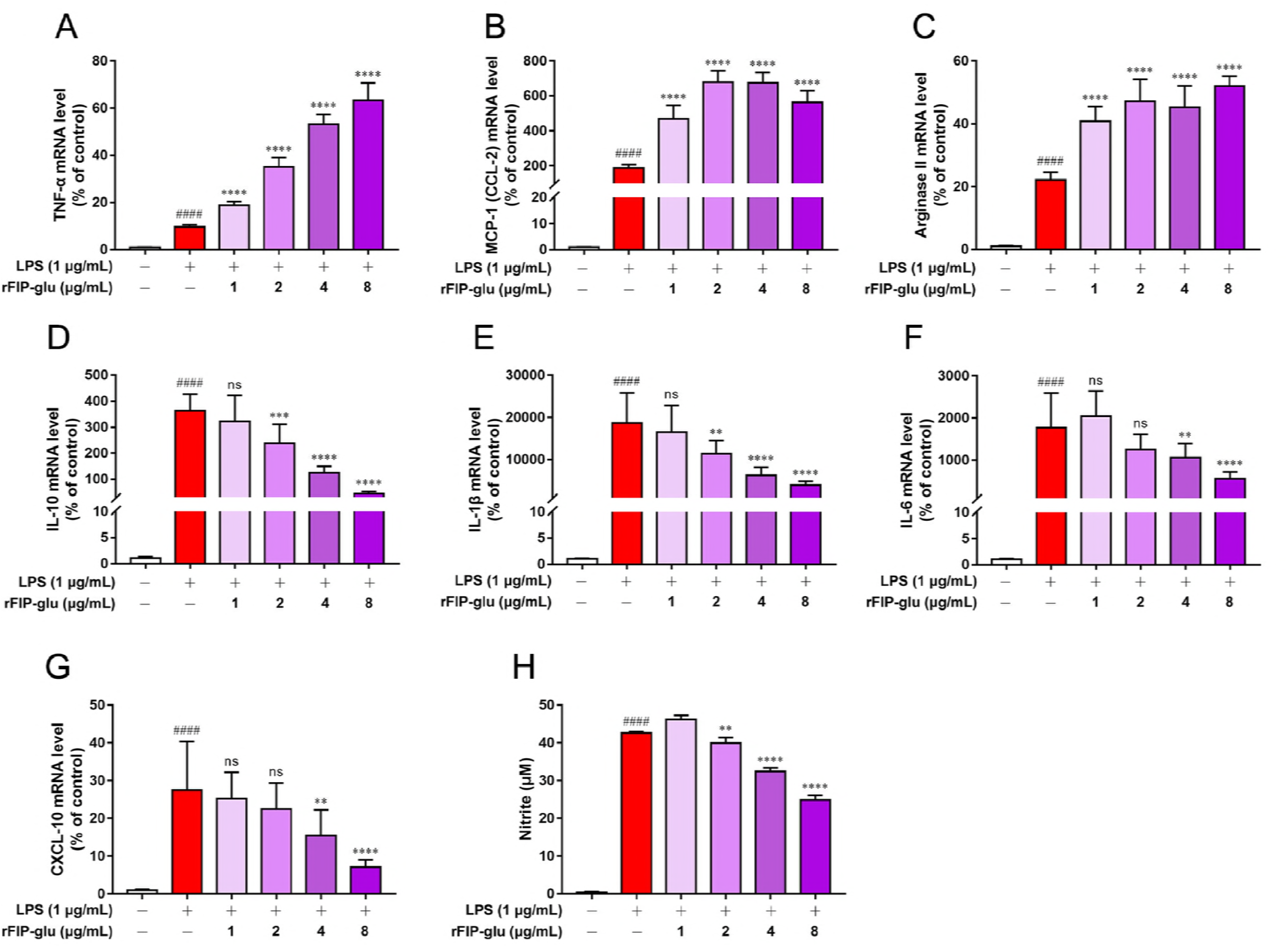
Effects of rFIP-glu on LPS-induced RAW264.7 cells. Cells were treated with different concentration of rFIP-glu (1, 2, 4 and 8 μg/mL) and LPS (1 μg/mL) and incubated for 6 h. The mRNA expression of TNF-α (A), MCP-1 (CCL-2) (B), Arginase II (C), IL-10 (D), IL-1β (E), IL-6 (F) and CXCL-10 (G) was measured by qRT-PCR. Data were expressed as means ± SD (n = 3). ####, p ≤ 0.0001 versus control group. ns, *p* > 0.05; **, *p* ≤ 0.01; ***, *p* ≤ 0.001; ****, *p* ≤ 0.0001 versus LPS group. The content of NO (H) in cell supernatant was determined by Griess assay. Values are means ± SD (n = 5). ####, *p* ≤ 0.0001 versus control group. **, *p* ≤ 0.01; ****, *p* ≤ 0.0001 versus LPS group.

### RNA sequencing results

To give further insights into the molecular mechanisms involved in the activity of rFIP-glu on macrophage RAW264.7 cells, RNA-seq was performed at the sequencing core facility of Shanghai Institute of Immunology. RAW264.7 cells were exposed to non-toxic rFIP-glu doses (4 μg/mL) for 6 h (Fig. 6). There were approximately 50,000 (coding and non-coding) genes detected. To determine sample relationships, principle component analysis (PCA) (Fig. 6A) and hierarchical clustering analysis (Heatmap) (Fig. 6B) were performed and demonstrated that two different groups, rFIP-glu treatment group and control (PBS) group, can be distinguished and showed segregation. Using an FDR ≤ 0.001 and fold change > 1 to determine differently expressed genes after rFIP-glu treatment, more than 700 genes were differentially expressed and we observed 578 up-regulated genes and 159 down-regulated gene (Fig. 6C). Next, we used the Gene Ontology (GO) database to characterize the differentially expressed genes. Subsets of these genes were found to be involved in cellular process, regulation of biological process, metabolic process, response to stimulus, developmental process, signaling and localization (Fig. 6D). These genes encoded proteins that perform immunological functions including inflammatory response, response to oxygen-containing compound, cellular response to organonitrogen compound, regulation of protein phosphorylation, regulation of protein modification process and regulation of programmed cell death (Fig. 6E). Moreover, to investigate the possible signaling pathways through which RAW264.7 cells activated by rFIP-glu, Kyoto Encyclopedia of Genes and Genomes (KEGG) pathway enrichment analysis was performed. Fig. 6F showed the top 20 significantly enriched canonical pathways. In order to verify the RNA-seq results, 10 differential expression genes which were up-or down-regulated showed in the RNA-seq results were selected. The expression levels of these genes measured by RT-qPCR showed the same tendency with RNA-seq (Fig. 6G).

**FIG 6.**
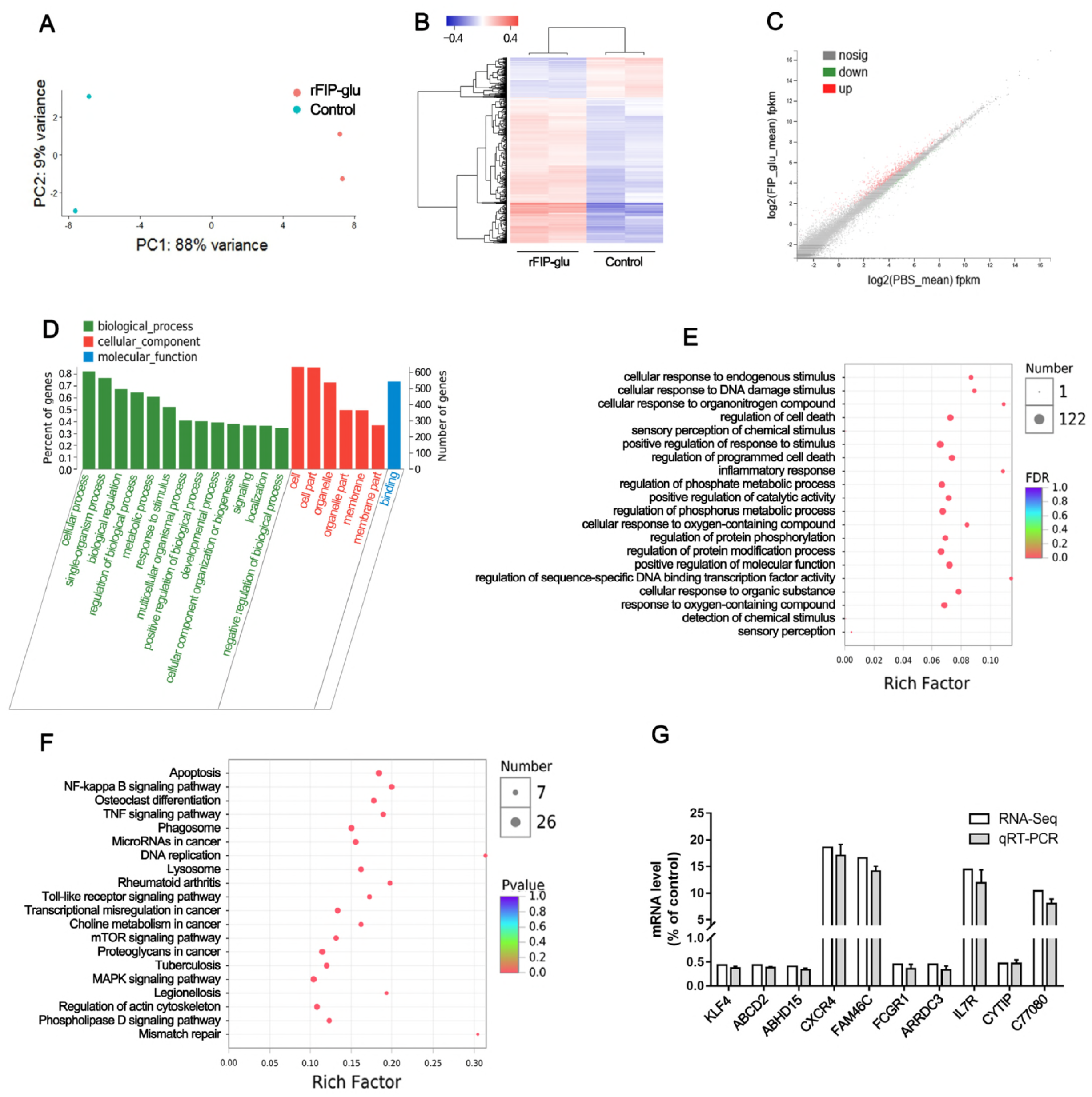
RNA-seq analysis of macrophage RAW264.7 cells. (A) Principle component analysis of RAW264.7 cells treated with and without rFIP-glu. (B) Heatmap of RAW264.7 cells treated with and without rFIP-glu. (C) Scatter plots of differentially expressed genes between PBS-and rFIP-glu-treated macrophages. Up-regulated genes are depicted in red and down-regulated gene in green (FDR ≤ 0.001, fold change > 1). (D) GO analysis of the differentially expressed genes. (E) GO enrichment analysis of the differentially expressed genes. (F) KEGG pathway enrichment analysis of the differentially expressed genes. (G) Confirmation of RNA-seq results by qRT-PCR.

### PI3K and MAPK signaling pathways are involved in rFIP-glu-induced macrophage activation

Obviously, results mentioned above had been shown that rFIP-glu was capable to promote macrophage proliferation and phagocytosis and induce the mRNA expression of inflammatory mediators such as TNF-α, MCP-1 (CCL-2) and iNOS. Analysis of RNA-seq indicated that Toll-like receptors pathway was involved in macrophage activation by rFIP-glu (Fig. S2). Signaling pathway inhibitor, PI3K inhibitor LY294002, was used to further confirm the mechanisms involved in rFIP-glu-induced macrophage activation. mRNA levels of TNF-α, MCP-1 (CCL-2) and iNOS were measure by qRT-PCR. Results showed that PI3K inhibitor LY294002 blocked the mRNA levels of MCP-1 (CCL-2) induced by rFIP-glu in RAW264.7 cells (Fig. 7B; *p* ≤ 0.0001), while the mRNA expression of TNF-α was not changed (Fig. 7A). These findings suggest that the PI3K pathway is related to rFIP-glu-induced macrophage activation.

**FIG 7.**
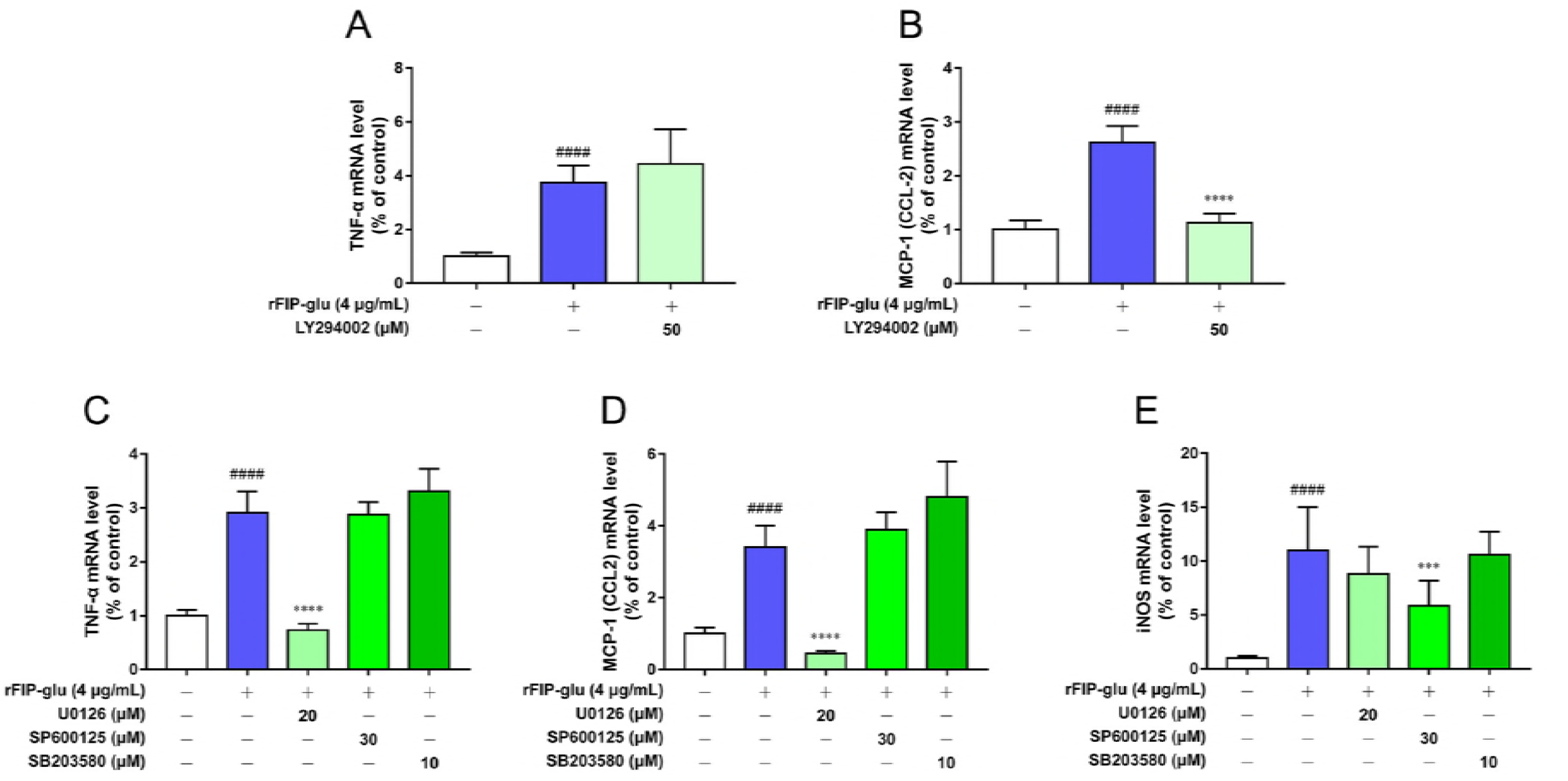
Effect of rFIP-glu on signaling pathways. RAW264.7 cells were pre-treated with 50 μM of LY294002 for 30 min and then treated with 4 μg/ml of rFIP-glu for 6 h. The mRNA expression of MCP-1 (CCL-2) (A) and TNF-α (B) was measured by qRT-PCR. RAW264.7 cells were pre-treated with 20 μM of U0126, 30 μM of SP600125 or 10 μM of SB203580 for 30 min and then treated with 4 μg/ml of rFIP-glu for 6 h. The mRNA expression of TNF-α (D), MCP-1 (CCL-2) (E) and iNOS (F) was measured by qRT-PCR. Data were expressed as means ± SD (n = 3). ####, *p* ≤ 0.0001 versus control group. ***, *p* ≤ 0.001; ****, *p* ≤ 0.0001 versus rFIP-glu group.

RNA-seq data implied that MAPK signaling pathway was also involved in macrophage activation by rFIP-glu (Fig. S3). Signaling pathway inhibitors, MEK1/2 inhibitor U0126, JNK inhibitor SP600125 and p38 inhibitor SB203580, were used to further confirm the mechanisms involved in rFIP-glu-induced macrophage activation. mRNA levels of TNF-α, MCP-1 (CCL-2) and iNOS were measure by qRT-PCR. Results showed that, in rFIP-glu-induced RAW264.7, MEK1/2 inhibitor U0126 blocked the mRNA levels of TNF-α (Fig. 7C; *p* ≤ 0.0001) and MCP-1 (CCL-2) (Fig. 7D; *p* ≤ 0.0001) and JNK inhibitor SP600125 prevented the up-regulation of iNOS mRNA (Fig. 7E; *p* ≤ 0.001). These findings suggest that the MEK1/2 and JNK pathways are indeed related to rFIP-glu-induced macrophage activation.

## Discussion

*G. lucidum* is well known as an edible medicinal mushroom for thousands of years in China. FIP-glu or LZ-8 is one of the active ingredients in *G. lucidum*. From beginning of discovery, FIP-glu is named immunomodulatory protein because of a certain degree of homology with the heavy chain variable region of several immunoglobulins (39, 40). FIP-glu possesses a variety of physiological activities, such as anti-anaphylaxis, proliferation stimulation of lymphocytes, anti-tumor and immunosuppression (7), suggesting that this protein has great potential in development of immuno-regulated drugs or foods. Our results showed that rFIP-glu is a potent stimulator of macrophage proliferation and activation. rFIP-glu can regulate the mRNA expression of pro-and anti-inflammatory mediators in macrophage RAW264.7 cells in the presence or absence of LPS. rFIP-glu mediates macrophage activation through PI3K and MAPK pathways based on RNA-seq analysis.

Macrophages play an important role in host-defense. Macrophages perform phagocytosis against pathogens as the first step, which is used to initiate the innate immune response, and then orchestrate the adaptive response (41). Thus, phagocytosis is a key indicator of evaluating macrophage activation. In the present study, rFIP-glu significantly improved the phagocytosis of macrophage RAW264.7 cells at 2 μg/mL, suggesting that rFIP-glu have abilities to enhance phagocytic activity of RAW 264.7 cells. The phagocytosis of macrophages can be enhanced by many bioactive substances, such as polysaccharides (42, 43), peptides (44) and proteins (45, 46), alkaloids (47) and phospholipids (48). Phagocytosis is one of the important innate immune responses. Following RNA-seq analysis showed that rFIP-glu-induced phagocytosis of macrophage RAW264.7 cells involved Fcγ receptor-mediated phagocytosis and most genes involved in this process were up-regulated (Fig. S4). There are two general classes of Fcγ receptors (FcγRs), activating receptors that activate effector functions and inhibitory receptors that inhibit these functions (41). In general, activation and inhibitory FcγRs are co-expressed on the same cell (49). The phagocytosis involves the simultaneous clustering of activating and inhibitory FcγRs and is regulated by the ratio of activating to inhibitory FcγRs (50). Moreover, FcγR-mediated phagocytosis is accompanied by the release of inflammatory mediators, and excessive magnitude of the FcγR response could lead to excessive inflammation (50, 51). RNA-seq analysis exhibited down-regulation of an activating receptor, FcγRI and up-regulation of an inhibitory receptor, FcγRIIb, when RAW264.7 cells were treated with 4 μg/mL of rFIP-glu although the enhancement of phagocytosis was not significant. This suggests that rFIP-glu participates in the regulation of the phagocytosis of macrophages and the release of inflammatory mediators. Additionally, FcγRIIb has ability to suppress allergic responses (52–55). This possibly is one of the mechanisms of rFIP-glu-mediated anti-allergy.

In response to an immune challenge, macrophages become activated and produce cytotoxic and inflammatory mediators, such as NO, ROS, TNF-α and IL-6, that contribute to nonspecific immunity (56). Our results implied that rFIP-glu has the predominant role in transcription of pro-inflammatory genes including TNF-α, MCP-1 (CCL-2), Arginase II and iNOS. Macrophages can be activated by biologically active substances such as polysaccharides and proteins to produce the pro-inflammatory molecules (24, 34, 57, 58). Similarly, most mushroom metabolites also activate macrophages to produce various mediators, such as IL-1β, TNF-α and iNOS (59). PCP, an immunomodulatory protein from *Poria cocos*, can promote TNF-α and IL-1β production in RAW 264.7 cells (60). TNF-α are pro-inflammatory cytokines and play an essential role in the immune response and inflammation (61). MCP-1 (CCL-2) is a member of the CC chemokine family and involved in the pathogenesis of multiple forms of inflammatory disorders as a mediator of acute and chronic inflammation (62, 63). Arginase II, one of isoforms of arginase, is up-regulated in M1 macrophages by pro-inflammatory stimuli and promotes pro-inflammatory responses (64, 65). iNOS-dependent nitric oxide from activated macrophages as a cytotoxic mediator can function in many diseases including cancer (66). Interestingly, in rFIP-glu-treated macrophage RAW264.7 cells, the mRNA expression of IL-6 was not changed, and that of IL-1β was not detected due to less transcripts possibly (Data not shown), although IL-1β and IL-6 are important pro-inflammatory cytokines as well. Moreover, the mRNA level of IL-10 was suppressed by rFIP-glu. IL-10 is an anti-inflammation cytokine and is secreted by M2 macrophages to suppress the inflammation (26, 67). Studies show that an increase in levels of M1 markers such as IL-1β, MCP-1 (CCL-2), TNF-α and iNOS and a decrease or little change in levels of M2 markers such as IL-10 will drive macrophage M1 activation (68–71). The findings of the present investigation contributed to our understanding that rFIP-glu promotes macrophage M1 polarization and initiates pro-inflammatory responses. Unexpectedly, CXCL-10 were down-regulated at mRNA levels in RAW264.7 cells induced by rFIP-glu. CXCL-10, belonging to the CXC family of chemokines, is involved in systemic inflammation and can mediate the recruitment of inflammatory cells (72, 73). Actually, CXCL-10 are expressed under inflammatory conditions in M1 macrophages (74, 75) and their products are reduced in M2 macrophages (76, 77). It seems that rFIP-glu induces M2 phenotypical macrophages to suppress inflammation. To further confirm anti-inflammatory effects of rFIP-glu, these related genes were detected in LPS-stimulated macrophage RAW264.7 cells. Results showed that rFIP-glu indeed possessed anti-inflammation activity through inhibiting LPS-induced mRNA levels of pro-inflammatory mediators (IL-6, IL-1β and CXCL-10) and the production of NO. A vast majority of bioactive substances can attenuate LPS-induced inflammation by decreasing the mRNA levels of pro-inflammatory mediators and increasing anti-inflammatory mediators. Polysaccharides, SGP-1 and SGP-2 isolated from the rhizomes of *Smilax glabra*, significantly suppressed the release of NO, TNF-α and IL-6 from LPS-induced RAW 264.7 cells (78). A prenylated flavonoid, 10-oxomornigrol F (OMF), can inhibit the LPS-induced production of NO, TNF-α, IL-1β and IL-6 in RAW264.7 cells (79). To our surprise, rFIP-glu acted in strong synergy with LPS to induce the mRNA expression levels of TNF-α, MCP-1 (CCL-2) and Arginase II. Meanwhile, mRNA level of IL-10 was still suppressed by rFIP-glu in LPS stimulated RAW264.7 cells. Another immunomodulatory protein, FIP-apo (APP) from *Auricularia polytricha*, also accounts for synergistic effects with LPS by NO and TNF-α production (80). These results mentioned above suggest that rFIP-glu can balance M1/M2 macrophages by regulating pro-and anti-inflammatory mediators and exhibit immunomodulatory activity. This phenomenon may confer macrophages an ability to quickly switch between M1 or M2 associated functions allowing for appropriate responses to stimuli and tissue environment (81). Re-polarization of macrophages is a key role and a promising therapeutic option in many diseases (26, 82). For example, inflammatory bowel diseases can be ameliorated by switching M1 macrophages to M2 (83, 84). Conversion of M2 to M1 phenotype is a potential therapeutic intervention in anti-tumor (85, 86).

Obviously, rFIP-glu has the ability to activate macrophages. The mechanisms were further investigated. In this study, RNA-seq was used to investigate the mechanisms of macrophage activation in rFIP-glu-treated RAW 264.7 cells. Our results indicated that the macrophage activation induced by rFIP-glu involved Toll-like receptors and MAPKs signaling pathways in macrophage RAW264.7 cells, which is consistent with that PI3K/Akt, the downstream of Toll-like receptor, and MAPK are involved in the activation of macrophages (34, 87–89). TLR4 is critical in immune responses and involved mainly in inflammation responses (90). TLR4 can be recognized and activated by many stimuli such as polysaccharides, and its expression increases, and then activates PI3K/Akt and MAPKs pathways, with introduction of a pool of inflammatory mediators (30, 34, 43, 87, 91). In addition, NF-κB involves in macrophage activation as well (24, 34, 92, 93). Similarly, RNA-seq analysis indicated that NF-κB pathway participated in rFIP-glu-mediated macrophage activation. More evidences should be investigated in the future. Although rFIP-glu activated Toll-like receptors, MAPKs and NF-κB pathways, RNA-seq (Fig. S2) and qRT-PCR (Fig. S5) confirmed that the mRNA expression level of TLR4 did not change. It suggests that the mRNA expression of pro-inflammatory genes induced by rFIP-glu was not through activating the TLR4 signaling pathway. RNA-seq analysis showed that TLR2 mRNA expression increased (Fig. S2), presuming TLR2 was a receptor of rFIP-glu. A preliminary yeast two-hybrid experiment showed that rFIP-glu did not interact with an extracellular part of TLR2 (Data not shown), suggesting TLR2 possibly was not a receptor of rFIP-glu as well. It is noteworthy that some active substances can activate all of these pathways (34, 57, 88), while some can only activate one or more (30, 87, 94). We next used specific PI3K/Akt and MAPKs pathway inhibitors to clarify whether these signaling pathways were involved in macrophage activation induced by rFIP-glu. Our results implied the involvement of PI3K in rFIP-glu mediated MCP-1 (CCL-2) mRNA production, MEK1/2 in TNF-α and MCP-1 (CCL-2), and JNK in iNOS. Although the induction of phosphorylated MAPKs and PI3K/Akt receptors was not evaluated by Western blot, the participation of them was confirmed by RNA-seq as well as inhibition of the effects induced by pretreatment with signaling pathway inhibitors. These results indicate that rFIP-glu may enter cells and act with MEK1/2, JNK or PI3K indirectly or directly, and another hypothesis is that rFIP-glu interacts with TLR2 within cells (Fig. 8). In addition, the synergistic activity of rFIP-glu and LPS implied that rFIP-glu could enhance the expression of downstream mediators that are generated by Toll-like receptors pathway (80). Heme oxygenase-1 (HO-1) is an anti-inflammatory enzyme and attenuates the inflammatory response (95), which can be regulated by nuclear factor erythroid 2-related factor 2 (Nrf2) in the inflammatory response (96). The Induction of HO-1 can be though MAPK and PI3K signaling pathways (94, 97, 98). In the present study, mRNA level of HO-1 was significantly increased in rFIP-glu-induced RAW264.7 cells (Fig. S6). This result implies that the anti-inflammation of rFIP-glu is possibly mediated by HO-1 in macrophage RAW264.7 cells.

**FIG 8.**
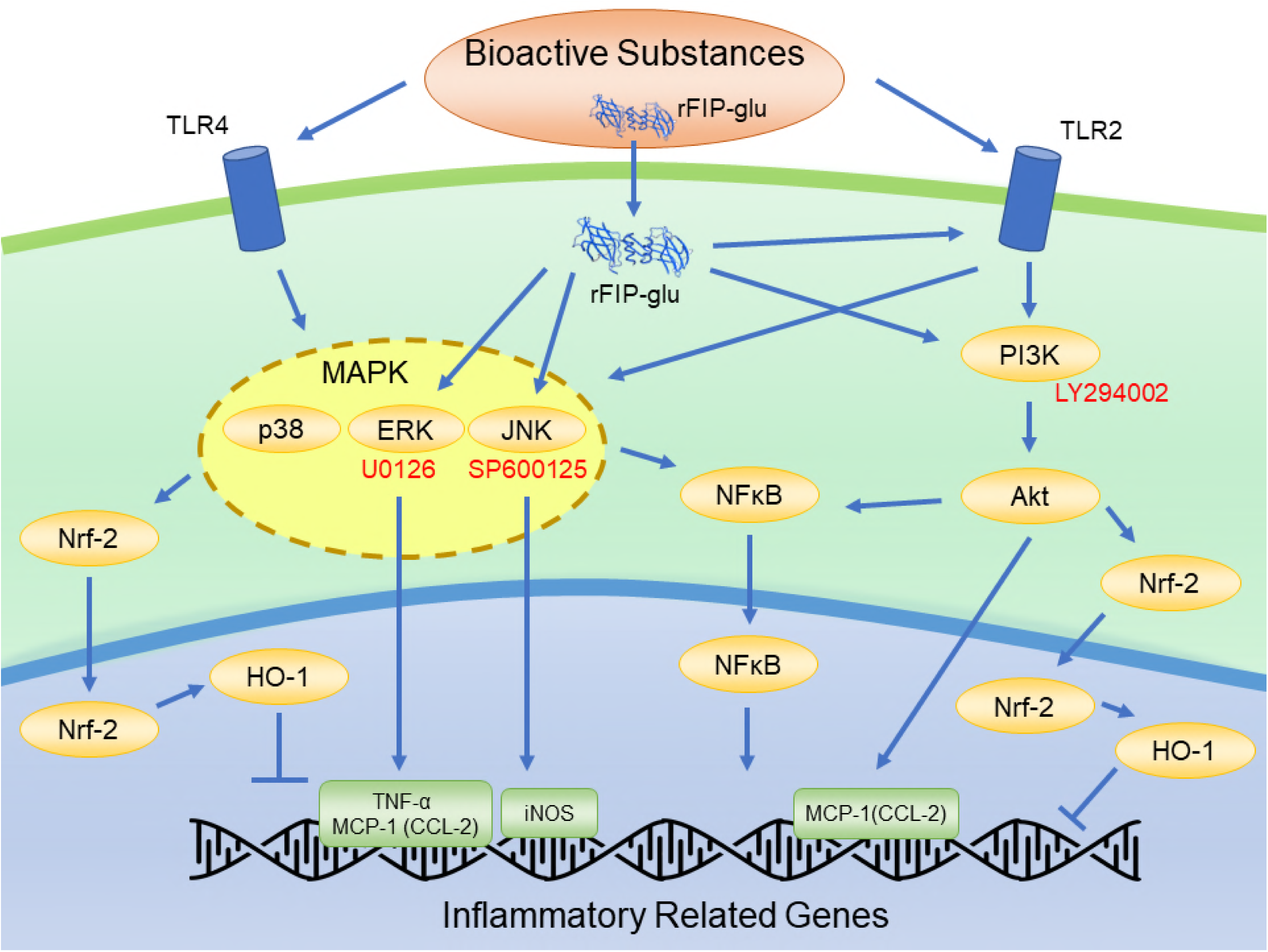
Possible immunomodulatory signaling mechanism of rFIP-glu in RAW264.7 macrophages.

## MATERIALS AND METHODS

### Reagents

Neutral red staining solution was obtained from Sangon Biotech (Shanghai, China). U0126 (MEK1/2 inhibitor), SP600125 (JNK inhibitor) and SB203580 (p38 inhibitor) were purchased from Beyotime (Shanghai, China). LY294002 (PI3K inhibitor) was purchased from Selleck Chemicals (Houston, TX, USA). Anti-FIP-glu antiserum was raised in rabbits (99). Anti-6×His Tag mouse monoclonal antibody, HRP-conjugated Goat Anti-Mouse IgG and HRP-conjugated Goat Anti-Rabbit IgG were from Sangon Biotech (Shanghai, China).

### Production of rFIP-glu

Production of recombinant FIP-glu (rFIP-glu) in *P. pastoris* was performed according to the instruction provided by Pichia Expression Kit (Invitrogen, USA). Briefly, a gene encoding FIP-glu and subcloned in pUC-57 vector was synthesized by Sangon Biotech (Shanghai) Co., Ltd. (China) based on codon usage bias. Then, the gene was cloned into an expression cassette vector pPIC9K. For convenience, a His-tag was added at 3’ end of multiple clone sites of the vector. After construction, the recombinant expression vector pPIC9K-glu-His linearized by a restriction enzyme *Sac* I was transferred into *P. pastoris* GS115. After confirmed by PCR (100) and sequencing, the transformant was induced by methanol for producing rFIP-glu. The rFIP-glu was purified with nickel-nitrilotriacetic acid (Ni-NTA) agarose resin (TaKaRa, Beijing, China). SDS-PAGE and Western blot analysis were performed based on the methods of our lab (99, 101).

### Cell culture

Macrophage RAW264.7 cells were purchased from the Cell Bank of the Chinese Academy of Sciences (Shanghai, China), cultivated in DMEM medium supplemented with antibiotics (100 U/mL penicillin and 100 mg/mL streptomycin) and 10% FBS and incubated at 37 °C in a 5% CO_2_ incubator. Cells were seeded at 2 × 10^5^ cells per well in 24-or 96-well microplates and were incubated in 5% CO_2_ at 37 °C.

### Cell viability

Methylene blue uptake assay was performed to assess the effect of rFIP-glu on the viability of macrophage RAW264.7 cells (102). Briefly, Raw264.7 cells were incubated for 24 h, and then were incubated with rFIP-glu at different concentration (1, 2, 4 and 8 μg/mL), LPS (1 μg/mL) and Concanavalin A (ConA) (5 μg/mL) for 24 h. LPS and ConA were used as positive controls. The culture supernatant was discarded and cells were stained by adding 50 μL of 0.6% methylene blue to each well. Plates were incubated at 37 °C for 60 min, and then inverted to drain the stain solution away. Wells were washed with phosphate buffered saline (PBS) to remove unbound stain. Plates were air-dried for several minutes. Stained cells were solubilized by adding 50 μL of Elution Buffer (Ethanol : PBS : Acetic acid = 50 : 49 : 1 (Volume)) for 20 min with a gentle shake. The absorbance value was measured at 570 nm using a microplate reader (BIO-TEK^®^, USA) and the viability was expressed as percentage versus control group.

### Phagocytosis assay

Phagocytosis assay was measured by Neutral red uptake assay (42). Briefly, Raw264.7 cells were incubated for 24 h, and then incubated with rFIP-glu at different concentration (1, 2, 4 and 8 μg/mL) and ConA (5 μg/mL) for 24 h. After the supernatant was discarded, 100 μL neutral red staining solution was added into each well and the plates were continued to incubate for 30 min. The supernatant was discarded and the cells were washed with PBS thrice to move free neutral red. 200 μL of cell lysis buffer (Ethanol : Acetic acid = 1 : 1 (Volume)) was added into each well and the plates were shaken for 2 h at room temperature. The absorbance value was measured at 540 nm using a microplate reader (BIO-TEK^®^, USA) and the phagocytosis was expressed as OD values.

### Measurement of NO

Raw264.7 cells were incubated for 24 h, and then were incubated with rFIP-glu at different concentration (1, 2, 4 and 8 μg/mL) and LPS (1 μg/mL) for 24 h. The supernatants were used to evaluate NO production using Griess assay by a NO Assay kit (Beyotime, Shanghai, China). According to the manufacturer’s protocol, sodium nitrite (NaNO_2_) was used to generate a standard curve to calculate the NO concentration.

### RNA extraction, sequencing and bioinformatics analysis

Macrophage RAW264.7 cells were cultured with rFIP-glu (4 μg/mL) for 6 h. Cells were harvested and total RNA was extracted using TaKaRa MiniBEST Universal RNA Extraction Kit (Beijing, China). Sequencing library was generated using Illumina Truseq stranded total RNA LT kit. On average 20 million Illumina paired-end reads (150 bp) were generated for each sample. The reads were mapped to reference using Hisat and differential expression analysis was performed using the R package DESeq2. In addition, principle component analysis (PCA) was carried out on the genes significantly expressed between all different groups. We combined results of two Enrichment analysis tools. One was derived from GSEA 3.0 desktop by mapping all genes to biological process of GO knowledge base and KEGG knowledge base. The other was calculated by IPA (version 01–13) of which cutoff was FDR ≤ 0.001 and Log2FoldChange > 1. The data were also analyzed on the free online platform of Majorbio I-Sanger Cloud Platform (www.i-sanger.com).

### cDNA synthesis and Real-time quantitative PCR (RT-qPCR)

cDNA was synthesized from 1 μg RNA using PrimeScript™ RT Master Mix (Perfect Real Time) (TaKaRa, Beijing, China) according to the manufacturer’s protocol. RT-qPCR was performed using SYBR qPCR Master Mix (Vazyme, Nanjing, China) and Roche LightCycler^®^96 Application. For PCR, samples were heated to 95 °C for 1 min, denatured at 95 °C for 20 s, annealed at 55 °C for 20 s, extended at 72 °C for 20 s, and cycled 45 times. The primers in this study were listed in Table S1. All reactions were performed in triplicate, and Ct values were normalized to β-actin. Relative expression was calculated using the 2 ^-ΔΔCt^ method.

### Statistical analysis

Graph Pad Prism 7 software was used to prepare graphs and statistical analysis. Data are expressed as means ± SD. The statistical analysis used One-way ANOVA analysis. Significance was indicated as ns, *p* > 0.05; *, *p* ≤ 0.05; **, *p* ≤ 0.01; ***, *p* ≤ 0.001; and ****, *p* ≤ 0.0001.

## ACKNOWLEDGMENTS

This study was funded by the Tibet Shenglong Industry Co., Ltd. (No. 2013310031001210).

## Supplementary legends

FIG S1 The effect of rFIP-glu on morphological changes of RAW264.7 cells. RAW264.6 cells were incubated with LPS, rFIP-glu or both for 6 h. Cells were subjected to microscopic analysis (200×). All experiments were performed in duplicates. The data from one representative experiment out of two independent experiments were shown. Bar is 50 μm. A, untreated cells. Untreated RAW 264.7 cells as control exhibited a round morphology. B, LPS-treated cells. LPS-treated RAW 264.7 cells exhibited changes of morphology including pseudopodia formation and cell spreading. C, rFIP-glu-treated cells. After stimulated by rFIP-glu, the volume of RAW 264.7 cells increased and agglutination happened. D, rFIP-glu and LPS-stimulated cells. rFIP-glu treatment reversed morphology change in LPS-stimulated RAW 264.7 cells, with increase in volume and agglutination.

FIG S2 Heatmap of Toll-like receptors pathway in RAW264.7 cells treated with and without rFIP-glu. RAW264.7 cells were treated with 4 μg/mL of rFIP-glu for 6 h.

FIG S3 Heatmap of MAPK receptors pathway in RAW264.7 cells treated with and without rFIP-glu. RAW264.7 cells were treated with 4 μg/mL of rFIP-glu for 6 h.

FIG S4 Heatmap of FcγR-mediated phagocytosis pathway in RAW264.7 cells treated with and without rFIP-glu. RAW264.7 cells were treated with 4 μg/mL of rFIP-glu for 6 h.

FIG S5 Effects of rFIP-glu on the mRNA level of TLR4 in RAW264.7 cells. The cells were treated with 4 μg/mL of rFIP-glu for 6 h. The mRNA expression of TLR4 was measured by qRT-PCR.

FIG S6 Effects of rFIP-glu on the mRNA level of HO-1 in RAW264.7 cells. The cells were treated with different concentrations of rFIP-glu (1, 2, 4 and 8 μg/mL) for 6 h. The mRNA expression of HO-1 was measured by qRT-PCR. Data were expressed as means ± SD (n = 3). ****, *p* ≤ 0.0001 versus control group.

TABLE S1 Primers used in this study.

